# NeuronID: An automatic toolkit for identifying neurons in two-photon calcium imaging data

**DOI:** 10.1101/2025.09.06.673975

**Authors:** Jikan Peng, Tian Xu

## Abstract

Two-photon calcium imaging has emerged as a powerful technique for monitoring neuronal activity in neuroscience; however, its data processing remains challenging. Here, we introduce NeuronID, an automatic toolkit designed to process two-photon calcium imaging data. The NeuronID toolkit features a modular architecture that includes motion correction, noise reduction, segmentation of neuronal components, and extraction of neuronal signals. Notably, the NeuronID toolkit offers an optimized strategy for segmenting neuronal components, which systematically integrates morphological boundary identification, cross-correlation analysis between pixels, and evaluation of neuronal signal quality. Compared to existing tools or manual annotation by experts, the NeuronID toolkit reduces the likelihood of over-segmentation while achieving near-human accuracy. Overall, this study provides a standardized analytical tool for processing two-photon calcium imaging data.

## Introduction

Monitoring single-neuron activity is essential for understanding the functional organization of the nervous system. Among available techniques for monitoring neuronal activity, two-photon calcium imaging has emerged as a powerful approach^1,2^. This approach detects neuronal activity by measuring changes in the fluorescence intensity of genetically encoded calcium indicators^3–7^. Two-photon calcium imaging has undergone substantial technological advancements, evolving from deep-tissue microscopy to high-speed volumetric imaging and miniaturized systems^1,8–11^. These developments now permit the recording of neuronal activity across entire brain regions in both head-fixed and freely moving animals. Importantly, the technique offers many advancements, including high-resolution visualization of neuronal morphology, precise anatomical mapping of single neurons, and detailed characterization of their spatiotemporal firing patterns over extended experimental periods.

However, as the scale and scope of two-photon calcium imaging continue to expand, the absence of standardized analytical tools has become a critical limitation. In particular, accurate segmentation of neuronal components (*e.g.*, soma) remains challenging, with existing methods often suffering from over-segmentation or suboptimal accuracy compared to manual annotation by experts^12–14^. To address this, we introduce NeuronID, an open-source, automatic toolkit for processing two-photon calcium imaging data. The NeuronID toolkit employs an optimized segmentation strategy that minimizes over-segmentation errors encountered in existing methods while achieving accuracy comparable to manual annotation by experts.

## RESULTS

### Overview of the NeuronID Toolkit

The NeuronID toolkit is an automatic, modular pipeline for two-photon calcium imaging analysis, including motion correction, noise reduction, segmentation of neuronal components, and extraction of neuronal signals. Its adaptable design supports continuous integration of novel algorithms, and a MATLAB implementation is publicly available (https://github.com/Peng-Jikan/NeuronID). The toolkit is deployed as a standalone desktop application (APP) with an intuitive graphical user interface (GUI), greatly enhancing its accessibility for experimental researchers without programming expertise (Figure 1A).

**Figure 1.**
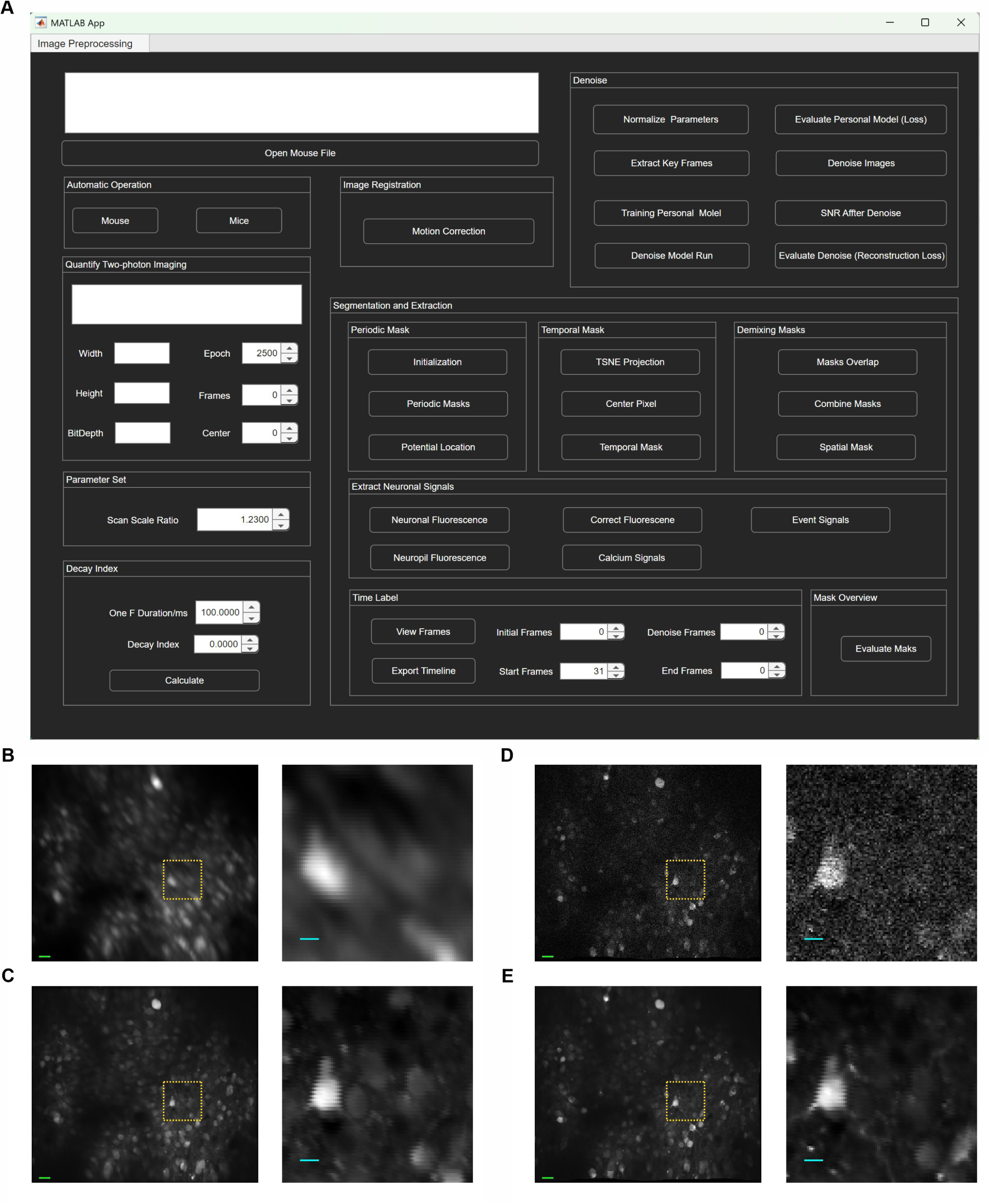
Motion correction and noise reduction. (A) The user-friendly GUI of the NeuronID toolkit, featuring an intuitive layout that guides users through the automated pipeline without requiring programming expertise. (B-C) Mean projection images before (B) and after (C) motion correction. (D-E) A typical frame before (D) and after (E) noise reduction. Scale bar: 10 pixels (green) or 30 pixels (blue).

### Motion Correction and Noise Reduction

Motion artifacts in two-photon calcium imaging data result from either rigid displacement or non-rigid deformations^15,16^. While multiple correction algorithms exist, we implemented the NoRMCorre algorithm in the NeuronID toolkit due to its proven efficacy and integration into widely used pipelines (Methods)^17^. This algorithm successfully corrected both types of motion artifacts, as indicated by improved sharpness in mean projection images before and after procession (Figure 1B and 1C). In addition, two-photon calcium imaging data is often contaminated by noise originating from electronic sources or the dynamics of calcium indicator^18^. To reduce this noise, we employed the DeepInterpolation algorithm, which utilizes an encoder-decoder deep network with skip connections to remove noise within each frame based on temporal context (Methods)^19^. This algorithm effectively reduced nearly all noise in individual frames (Figure 1D and 1E).

### Asynchronous Firing Gives Rise to Segmentation of Neuronal Components

In the NeuronID toolkit, we utilized asynchronous firing to segment neuronal components in two-photon calcium imaging data. First, we observed asynchronous firing across neurons, where only a subset of all recorded neurons was active in any given imaging frame (Figure 2A). This characteristic allowed us to identify distinct subsets of neuronal components across different time periods and then combine them to segment all neuronal components (Figure 2A). To implement this strategy, we divided the recording into temporally continuous periodic blocks (Figure 2B; Methods). Second, we discovered asynchronous firing between central pixels and boundary pixels within individual neurons. Specifically, central pixels maintained stable fluorescence signals during active periods, while boundary pixels exhibited transient intensity changes, leading to significantly greater max-mean intensity differences at boundary (Figure 2C, 2D and 2E). This characteristic allowed us to highlight the morphological boundaries of neuronal components in each periodic block by using the corresponding max-mean projection image (Figure 2F; Methods). To identify the morphological boundaries of neuronal components in each periodic block, we normalized the max-mean projection image and applied an adaptive threshold filter to generate periodic masks containing distinct regions of interest (ROIs) (Figure 2G). Subsequently, we refined the ROIs in the periodic masks using morphological operations (closing enclosed voids) and area criteria (by default, 5μm in diameter) (Figure 2G). Finally, each ROI in the periodic masks was assigned a unique identifier, with all pixels belonging to the same ROI sharing the same ID.

**Figure 2.**
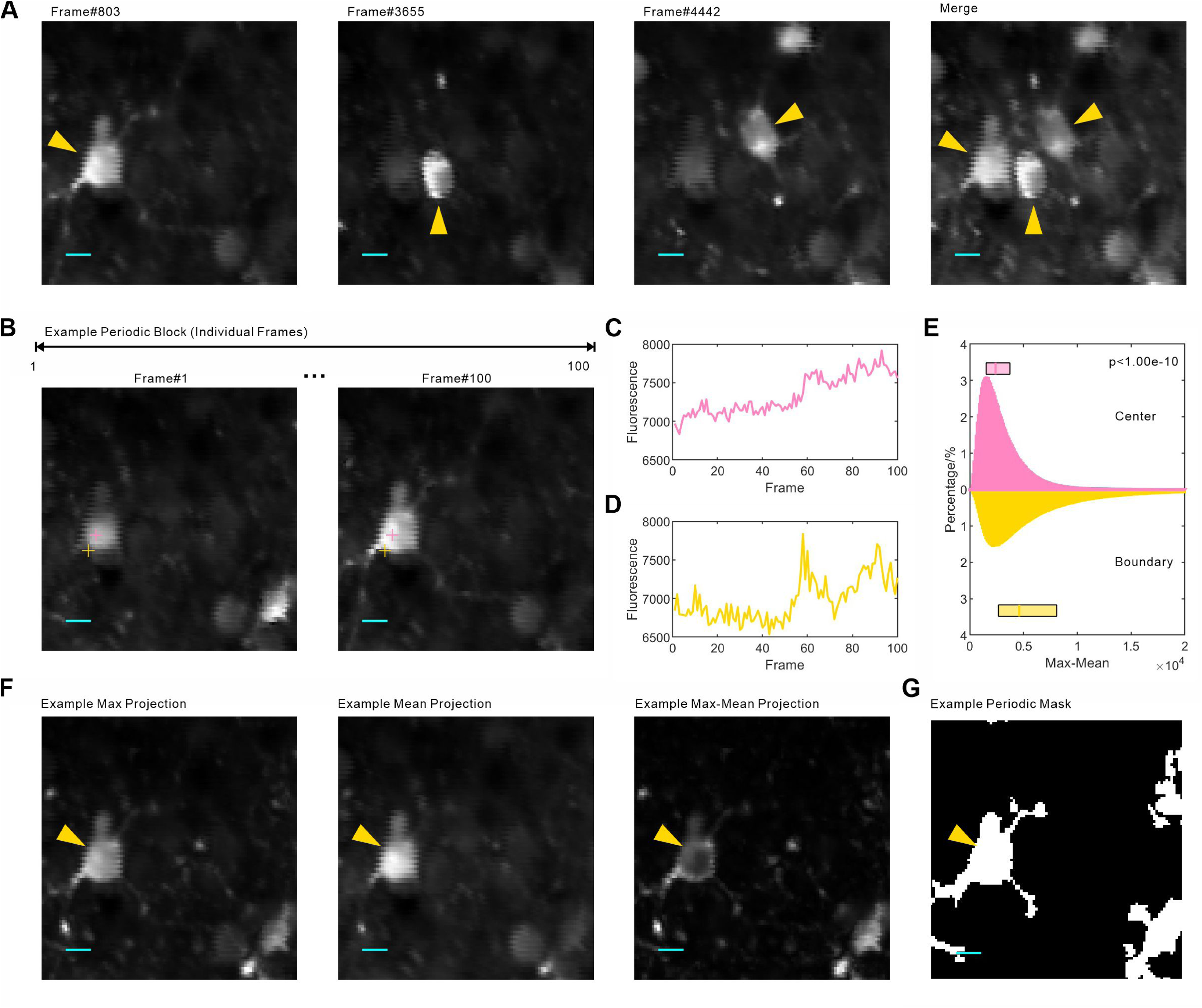
Asynchronous firing facilitates the segmentation of neuronal components. (A) Three representative neuronal components at different frames and in the corresponding merged image. (B) An example of a periodic block, with the positions of two representative pixels indicated. (C, D) Fluorescence signals of two labeled pixels during the periodic block. (E) Distribution of the max-mean values of central pixels or boundary pixels within a given neuronal component. Mann–Whitney U test. (F) Example images of maximum projection, mean projection, and max-mean projection. (G) A representative periodic mask. Scale bar: 30 pixels.

### Cross-correlation Analysis Refines Segmentation of Neuronal Components

As previous described, integrating ROIs across periodic masks is necessary to segment all neuronal components in two-photon calcium imaging data. To achieve this, we first retained pixels detected as an ROI in at least three distinct blocks, while classifying those with fewer occurrences as background (Figure 3A). We then grouped pixels sharing highly similar ID sequences with a Jaccard Index (JI) threshold of 0.5 (Figure 3B and 3C; Methods). However, performing cross-correlation analysis on all pixel pair imposed a substantial computational burden. For instance, in our example dataset, background segmentation left 280,338 pixels, resulting in over 39.3billion pixel pairs. To mitigate this, we identified 9,000 key representative pixels using an unsupervised clustering method and computed JI values only between these key pixels and their 1,000 neighboring pixels (Figure 3D and 3E; Methods). Consequently, we integrated periodic masks into a temporal mask (Figure 3F). In the temporal mask, we observed that some ROIs shared pixels, which could result from repeated segmentation of the same neuronal component or overlapping field of view among distinct neuronal components (Figure 3G). To address this, we merged any two overlapping ROIs if more than 80% of their pixel pairs met the JI threshold. Finally, a spatial mask was derived from the temporal mask (Figure 3H).

**Figure 3.**
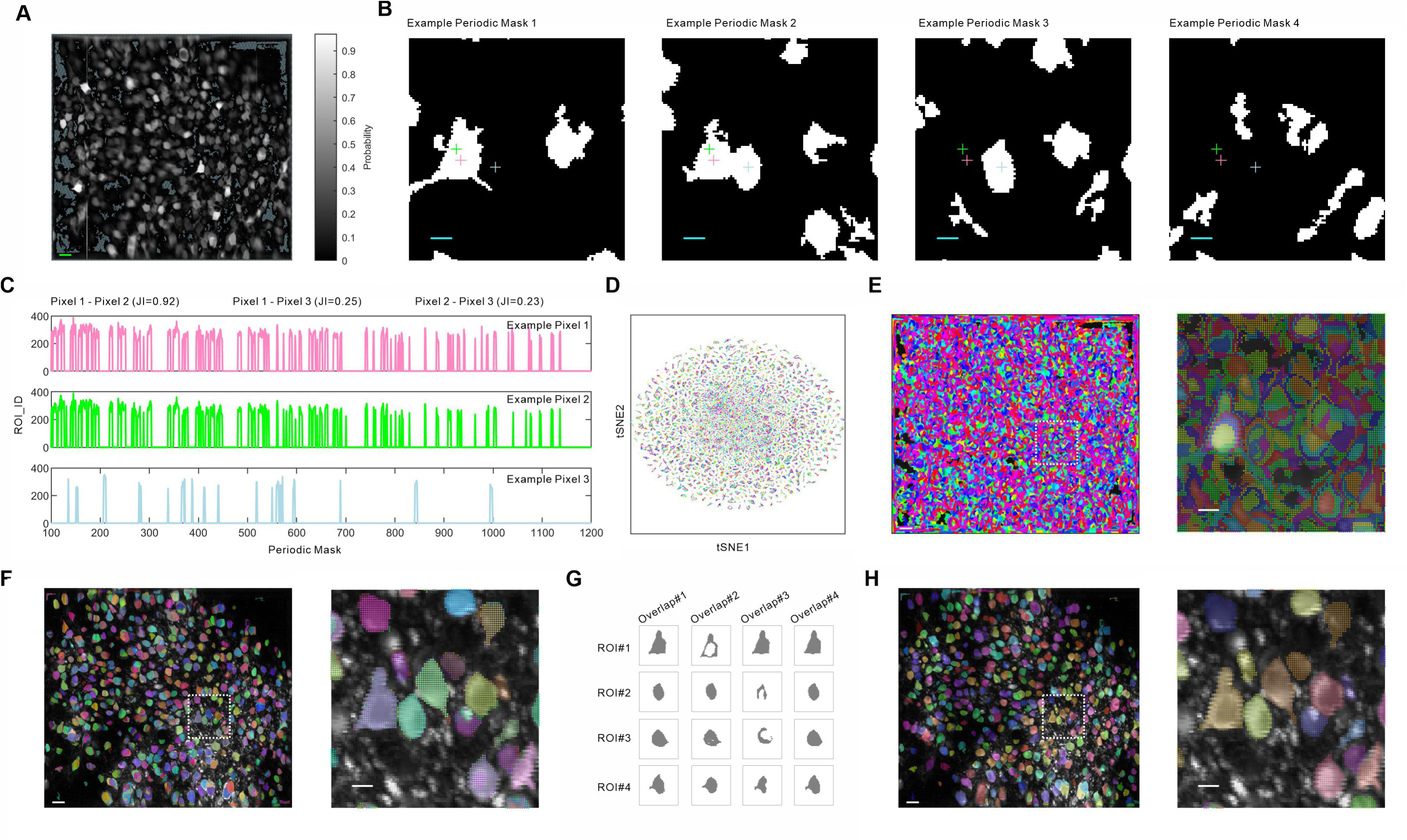
Segmentation of background and neuronal components. (A) Probability distribution for pixel classification into ROIs and segmentation of background (light green). (B) Four representative periodic masks with the positions of three selected pixels indicated. (C) ID sequences of the three pixels across periodic blocks. (D) t-SNE visualization of individual pixels, revealing 9,000 clusters identified by K-means clustering. (E) Distribution of the 9,000 clusters within the max-mean projection image. (F) Temporal mask. (G) Four representative ROIs repeatedly segmented. (H) Spatial mask. Scale bar: 10 pixels in A and the left panels of E, F, and H; 30 pixels in B and the right panels of E, F, and H.

### Extraction of Neuronal Signals

For each ROI in the spatial mask, we first identified its constituent pixels and selected an equivalent number of surrounding background pixels as neuropil (Figure 4A). We then extracted the fluorescence signals of both the ROI and neuropil by calculating the mean fluorescence intensity of their respective pixels in each frame (Figure 4B, Figure S1A and 1B). ROIs were retained if their fluorescence signals exceeded the 3σ level of their corresponding neuropil signals for at least three consecutive frames (Figure 4B). Next, we refined the ROI fluorescence signals by reducing neuropil contamination using a gradient reduction algorithm and removing photobleaching artifacts from the calcium indicators by eliminating polynomial trends in fluorescence decay over time (Figure 4C). Subsequently, we extracted the calcium signal of the ROI from the refined fluorescence signal using the ΔF/F metric, where ΔF represents the deviation from baseline fluorescence (F), estimated as the mode fluorescence intensity (Figure 4C, 4D and Figure S1C).

**Figure 4.**
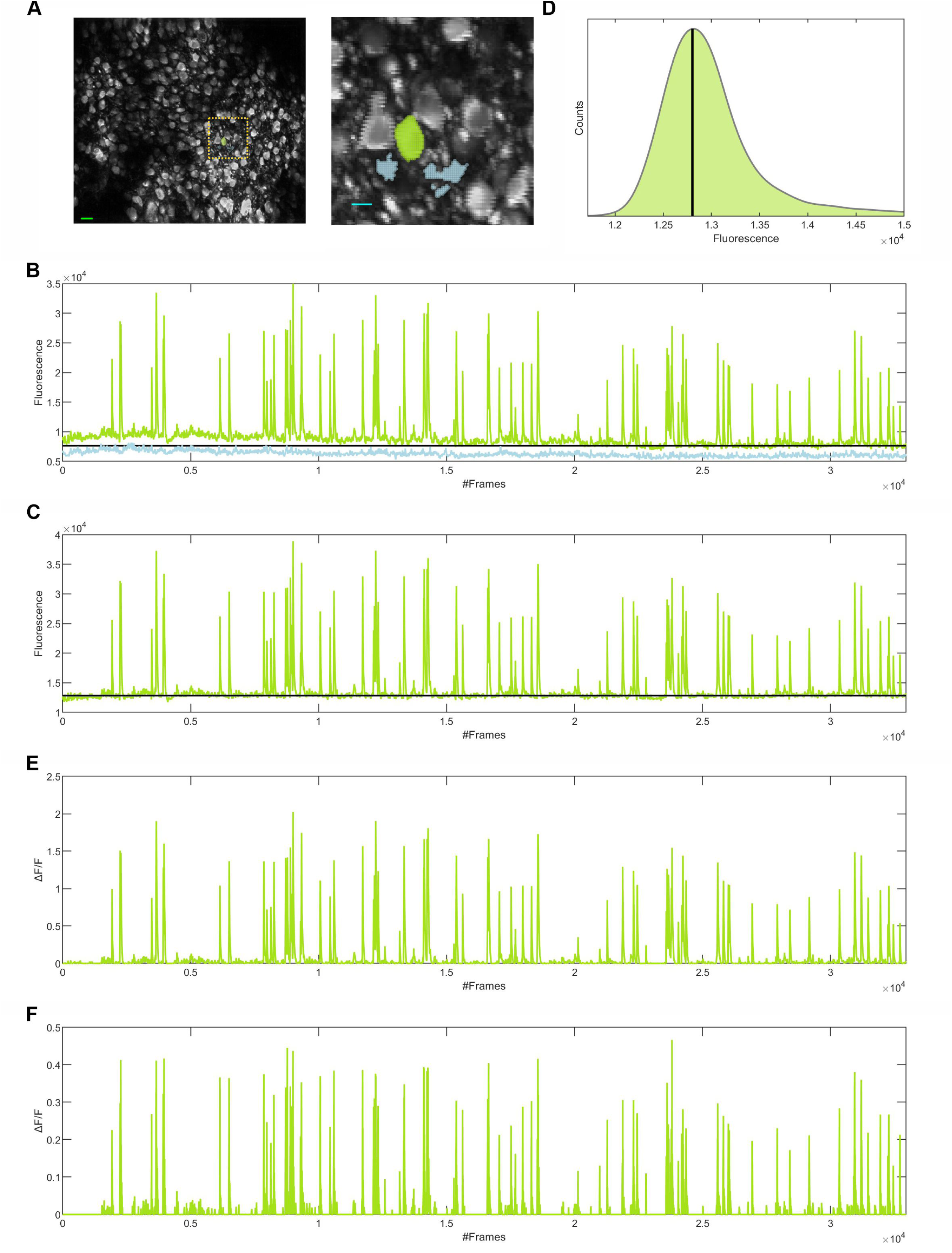
Extraction of neuronal signals. (A) The illustration of a ROI (green) and its corresponding neuropil region (blue). Scale bar: 10 pixels (green) or 30 pixels (blue). (B) Fluorescence signals of the ROI (green) and the neuropil (blue) across individual frames. The black line indicates the 3σ level of the neuropil fluorescence signal. (C) Fluorescence signal of the ROI after neuropil contamination reduction and polynomial trend removal. The black line represents the baseline level. (D) Distribution of fluorescence intensities, with the baseline indicated. (E) Calcium signal. (F) Event signal.

We further selected ROIs as neuronal components if they exhibited a ratio of positive-to-negative transients greater than ten (Methods) and removed the negative calcium signals to isolate calcium signal of the neuronal component (Figure 4E and Figure S1C). Finally, we extracted event signal of the neuronal component by deconvolving its calcium signal using imaging parameters (100ms frame rate, 860ms decay constant of calcium indicator) and a 2σ threshold of the calcium signal (Methods)^20^. The resulting positive signal within the event signal represented the firing status of the neuronal component, while the remaining signal represented the resting status (Figure 4F).

### Accuracy of the NeuronID Toolkit

Following the identification of morphological boundary, cross-correlation analysis between pixels, and evaluation of signals, the NeuronID toolkit successfully performed the segmentation of neuronal components, as illustrated in the soma mask (Figure 5A, Figure S2, and Figure S3; Methods). To evaluate its accuracy, we compared it against existing tools (CaImAn and Suite2p) or manual annotation by experts^12–14^. First, we observed that some neuronal components were over-segmented in existing tools (Figure 5A and Figure S3). We further observed that the morphological boundaries of some neuronal components were recognized more accurately in the NeuronID toolkit (Figure 5B and 5C). These observations implied that the NeuronID could reduce the likelihood of over-segmentation. Indeed, the number of neuronal components segmented by the NeuronID toolkit was significantly fewer than that segmented by the two existing tools (Figure 5D; Table S1). Such reduction in over-segmentation in the NeuronID toolkit could result from the identification of morphological boundary, which was not employed by CaImAn and Suite2p. Furthermore, the NeuronID toolkit conducted cross-correlation analysis between pixels across periodic blocks rather than individual frames, which increased the likelihood of identifying pixels belonging to the same neuronal components (Figure 5E).

**Figure 5.**
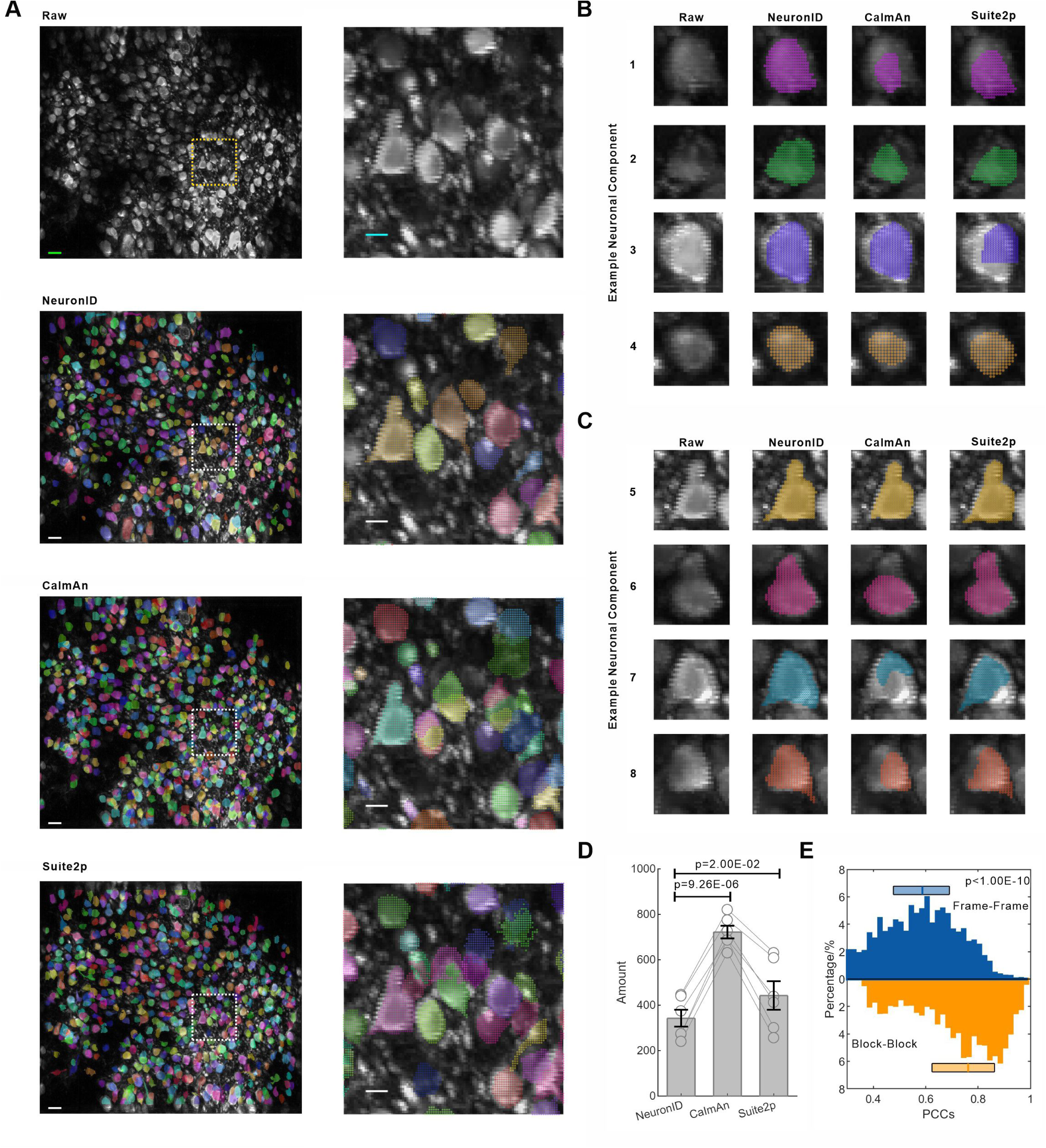
Comparative analysis of the NeuronID toolkit against two existing tools. (A) Max-mean projection image (Raw) and soma masks generated by NeuronID, CaImAn, and Suite2p. Scale bar: 10 pixels (left) or 30 pixels (right). (B) Representative examples of somatic structures segmented by the three toolkits. (C) Examples of complex neuronal components segmented by the three toolkits. (D) Number of neuronal components segmented by the three toolkits. Paired t-test. (E) Distribution of Pearson correlation coefficients (PCCs) between pixels within a given neuronal component across individual frames or periodic blocks. Mann-Whitney U test.

Second, we observed that segmentation of neuronal components using the NeuronID toolkit showed high concordance with manual annotation by experts (Figure 6A and Figure S4). We further observed that the identification of morphological boundaries in some neuronal components approached the accuracy of manual annotation (Figure 6B). Our analysis revealed that over 80% of neuronal components were accurately segmented by the NeuronID toolkit, as measured by overlap ratio (Figure 6C; Methods). The NeuronID toolkit achieved an average F1 score of 0.87, indicating near-human performance (Figure 6D; Table 1). Together, these results established the NeuronID toolkit as a robust solution for reducing over-segmentation while maintaining high segmentation accuracy.

**Figure 6.**
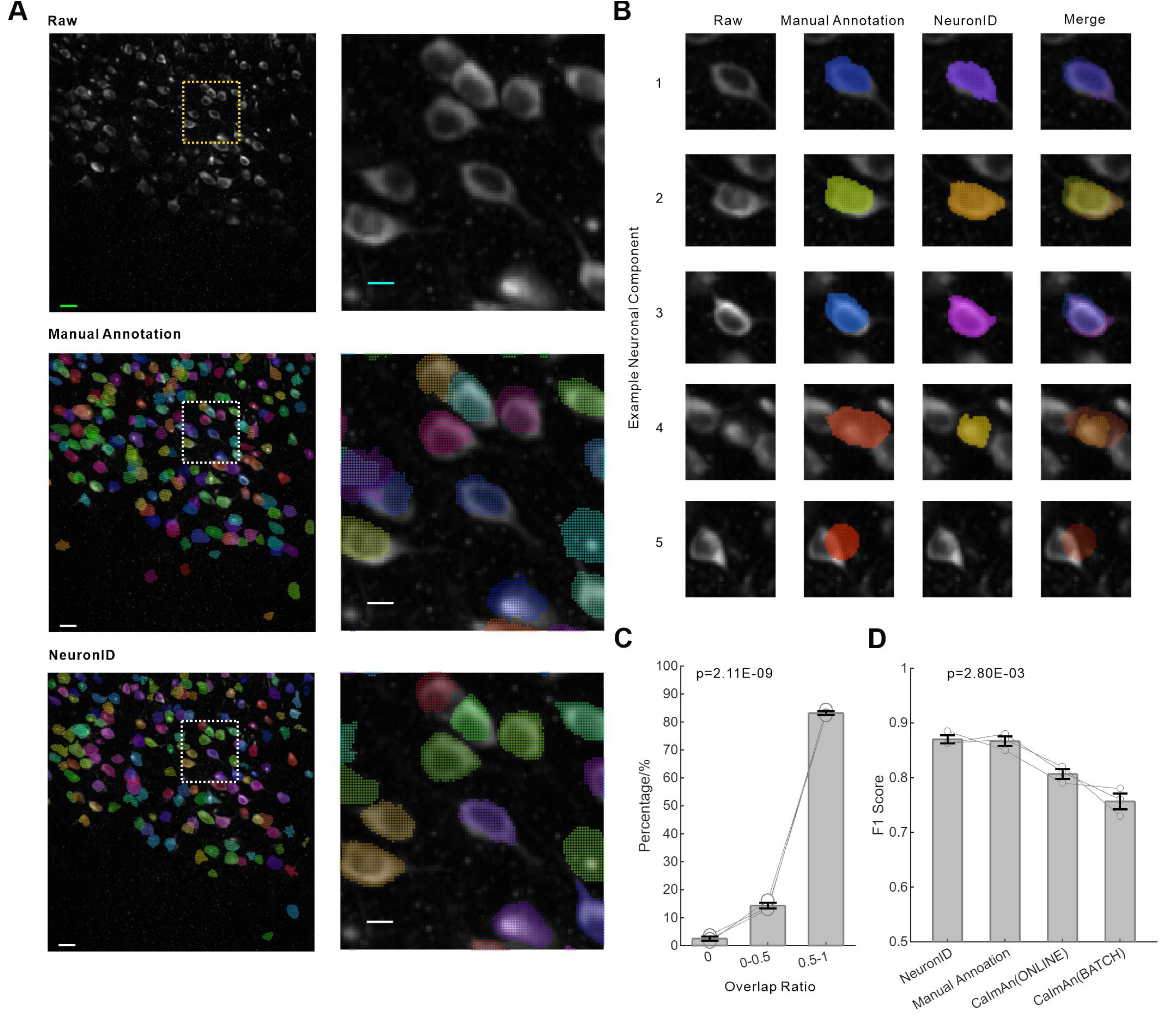
Comparative analysis of the NeuronID toolkit versus manual annotation by experts. (A) Max-mean projection image (Raw) and soma masks generated by manual annotation or the NeuronID toolkit. Scale bars: 10 pixels (left) and 30 pixels (right). (B) Representative examples of neuronal components segmented by both methods. (C) Distribution of the overlap ratio. One-way ANOVA. (D) F1 score of different methods. One-way ANOVA.

**Table 1.** Accuracy of different methods.

| Dataset ID | CalmAn (ONLINE) <sup>a</sup> | CalmAn (BATCH) <sup>a</sup> | Manual Annotation <sup>b</sup> | NeuronID <sup>c</sup> |
| --- | --- | --- | --- | --- |
| J123 | 0.82, 172 | 0.73, 157 | 0.87, 183 | 0.86, 160 (148, 12, 35) |
| J115 | 0.79, 1091 | 0.78, 738 | 0.85, 891 | 0.86, 794 (727, 67, 164) |
| K53 | 0.81, 1025 | 0.76, 809 | 0.88, 920 | 0.89, 816 (769, 47, 151) |

<sup>a</sup>F1 score against manual annotation by experts<sup>14</sup>
<sup>b</sup>F1 score against different experts<sup>14</sup>
<sup>c</sup>F1 score against manual annotation by experts, with the values of TP, FP, FN.

## Discussion

In this study, we introduced the NeuronID toolkit, which provides a standardized and automated pipeline for analyzing two-photon calcium imaging data. The NeuronID toolkit integrated four core modules, including motion correction, noise reduction, segmentation of neuronal components, and extraction of neuronal signals. Notably, the NeuronID toolkit offered an optimized strategy for segmenting neuronal components, which systematically combines morphological boundary identification, cross-correlation analysis between pixels, and evaluation of neuronal signal. Compared to existing tools that rely solely on cross-correlation analysis or manual annotation focusing on morphology^12–14^, the NeuronID toolkit reduced the likelihood of over-segmentation while achieving accuracy comparable to that of human experts.

However, the current version of NeuronID is specifically designed for two-photon calcium imaging data capturing the somatic structures of hundreds of neurons. Future developments should aim to extend its capabilities to process complex subcellular structures (*e.g.*, axons, dendrites, and synapses) as well as to handle volumetric data^21–24^. The modular architecture of the NeuronID toolkit is intentionally designed to support future enhancements and adaptability.

## Materials and Methods

### Two-photon datasets collection

In this study, we utilized six two-photon calcium imaging datasets from a previous study (Example Datasets 1-6)^25^. In these datasets, neuronal activity in layer II/III of the Posterior Parietal Cortex (PPC) was recorded in freely moving mice using FHIRM-TPM microscopy at a resolution of 600×512 pixels (487.80×416.20 μm) and a frame rate of 10 Hz for 60 minute. For validation, we used three additional expert-annotated benchmark datasets from CaImAn (J115, J123, and K53)^14^.

### Motion correction

In the NeuronID toolkit, we utilized the NoRMCorre algorithm to correct both rigid translations and non-rigid deformations in two-photon calcium imaging data^17^. To generate an initial reference template, we uniformly sampled 60 frames across the recording (one frame every 500 frames), ensuring coverage of diverse neural activity states. Notably, we implemented the latest Python version of NoRMCorre, while the original GitHub repository maintains only MATLAB code.

### Noise reduction

In the NeuronID, we incorporated the DeepInterpolation algorithm to reduce noise in two-photon calcium imaging data^19^. This approach utilizes an encoder-decoder deep network architecture with skip connections, which reconstructs each target frame from its temporal context (30 preceding frames and 30 succeeding frame). Given that calcium indicator dynamics are scanning-frequency dependent, we provided multiple pre-trained DeepInterpolation networks optimized for different acquisition rates. For further customization, the NeuronID toolkit included a transfer learning module, enabling users to finetune models to their specific imaging conditions. By default, this module selects 360 representative frames (spaced at 100-frame intervals) and performs 10,000 training iterations, though these parameters remain user-adjustable. To ensure optimal performance, the NeuronID toolkit offers two key evaluation metrics: (1) reconstruction loss, calculated as the mean absolute difference in fluorescence intensity between original and reconstructed frames, and (2) pixel-wise signal-to-noise ratio (SNR), derived from the ratio of mean intensity to standard deviation for each pixel across all frames.

### Max projection, mean projection, and max-mean projection

By default, each periodic block comprises 100 sequential frames, with a 20-frame gap between adjacent blocks. For each periodic block, we converted all sequential frames into mean or max projection image. Specifically, the mean projection image was generated by calculating the average fluorescence intensity of individual pixels across all frames. The max projection image was generated by determining the highest fluorescence intensity of individual pixel across all frames. Additionally, the max-mean projection image was generated by comparing these two projection images.

### Identification of key pixels and calculation of Jaccard Index

For each pixel retained after background segmentation, we constructed a feature vector comprising its spatial coordinates, the probability of being classified as an ROI, and the ID sequence across periodic masks. We reduced the dimensionality of these features using Principal Component Analysis (PCA), retaining 90 principal components (covering 95% variance)^26^. Next, we projected the reduced data into a 2D embedding space using t-SNE^27^. To group pixels, we applied K-means clustering to the t-SNE embedding, selecting 9,000 clusters based on the size and number of neuronal components^28^. The cluster centers were designated as key pixels, with their 1,000 nearest neighbors in t-SNE space defining associated neighboring pixels. To quantity similarity between two pixels, we computed the Jaccard Index as the ratio of periodic masks where both pixels shared an ROI assignment to the total number of masks where either pixel was assigned to an ROI.

### Refinement of fluorescence signal and identification of calcium transient

In the NeuronID toolkit, we implemented a gradient reduction algorithm to reduce neuropil contamination in ROI fluorescence signals^12,29^. This algorithm iteratively adjusted the neuropil signal using a correction factor (ranging from 0 to 1) to minimize the residual difference between the raw ROI fluorescence signal and the neuropil-subtracted signal^12^. Furthermore, we estimated and removed a polynomial trend from the fluorescence signal to account for photobleaching of calcium indicators, which is characterized by a gradual decline in fluorescence intensity over the course of imaging^6,30^. Additionally, we applied a 2σ threshold-based method to detect positive or negative transients in the calcium signal^31^. Specifically, positive transients were defined as increases in the calcium signal exceeding +2σ, where σ was calculated from the negative calcium signal; negative transients were defined as decreases exceeding -2σ, with σ derived from the positive calcium signal.

### Comparative analysis

To evaluate the accuracy of the NeuronID toolkit, we compared its performance to CaImAn and Suite2p^13,14^. We applied these tools to six example datasets, using their recommended default parameters. For each neuronal component segmented in the NeuronID toolkit, we computed the Pearson correlation coefficient (PCC) between pixels based on either fluorescence signals across frames or ID sequences across blocks. To further validate the NeuronID toolkit, we compared its performance with manual annotation by experts using three additional datasets^14^. Each dataset was independently labeled by two to three experts, and consensus masks were used as the ground truth. We assessed the overlap between neuronal components identified by the NeuronID toolkit and those labeled by experts. For each neuronal component labeled by experts, we calculated the overlap ratio with each neuronal component segmented in the NeuronID toolkit, selection the maximum value as the best match. Based on the distribution of overlap ratios, we calculated true positives (TP, the number of neuronal components with a maximum overlap ratio exceeding 0.5), false positives (FP, the number of neuronal components with a maximum overlap ranging from 0 to 0.5), and false negatives (FN, the number of neuronal components detected by experts but missed by the NeuronID toolkit)^32^. From these values, we derived precision (TP/(TP+FP)), recall (TP/(TP+FN)), and the F1score (harmonic mean of precision and recall) (Table 1)^32^.

### Quantification and statistical analysis

In this study, we employed various statistical methods to analyze the data, including the paired t-test, the one-way ANOVA, and the Mann-Whitney U test. P-values from the Mann-Whitney U test were adjusted using the Benjamini-Hochberg method. P-values were reported using scientific notation and rounded to two decimal places. In our data presentation, we utilized a variety of graphical representations to effectively convey our findings. Specifically, bar graphs were employed to illustrate mean values accompanied by Standard Error of the Mean (SEM) as error bars, with individual data points represented as dots for clarity (e.g., Figure 5D). Moreover, a specific graph was used to illustrate the distribution of the original data, presented through median values and interquartile ranges (e.g., Figure 2E).

## RESOURCE AVAILABILITY

### Lead contact

Requests for further information and resources should be directed to and will be fulfilled by the lead contact, Tian Xu

### Materials availability

Correspondence and requests for materials should be addressed to the lead contact.

### Data and code availability

All data are available in this manuscript. Codes for the NeuronID toolkit are available from the authors on request or the GitHub platform.

## Acknowledgments

We thank Hongyan Yang for technical assistance and Xu lab members for discussion. This study was supported in part by the grants to TX including National Natural Science Foundation of China (U21A20201), Key Laboratory of Growth Regulation and Translational Research of Zhejiang Province (2020E10027), the Science Technology Department of Zhejiang Province (2021ZY1019, 2022ZY1005), Zhejiang Leading Innovative and Entrepreneur Team Introduction Program (2018R01003), and Westlake Laboratory of Life Sciences and Biomedicine (202208011). JKP are supported by Westlake University Predoctoral Fellowship. TX is grateful to Yale and HHMI for more than two decades of support.

## Author contributions

Conceptualization, JKP and TX; Methodology, JKP; Investigation, JKP; Visualization, JKP; Funding acquisition, TX; Project administration, TX; Supervision, TX; Writing – original draft, JKP and TX; Writing – review & editing, JKP and TX.

---

**Figure S1.**
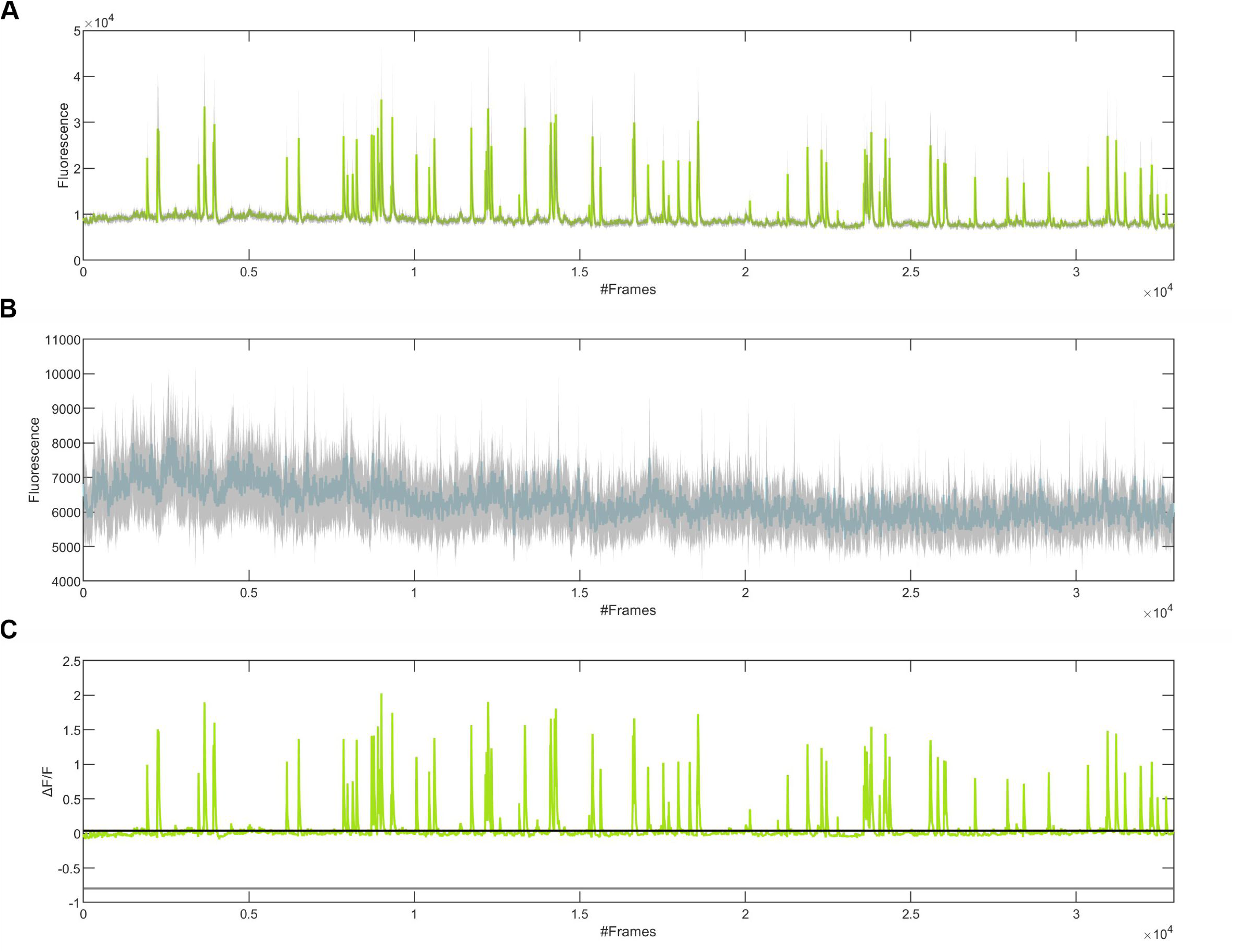
Signals of a ROI and its neuropil. (A, B) Average fluorescence intensity (green or blue) and corresponding σ values (gray) across individual frames for a ROI (A) and its neuropil (B). (C) Calcium signal of the ROI, with a +2σ threshold (black) indicating positive transients and a -2σ threshold (gray) indicating negative transients.

**Figure S2.**
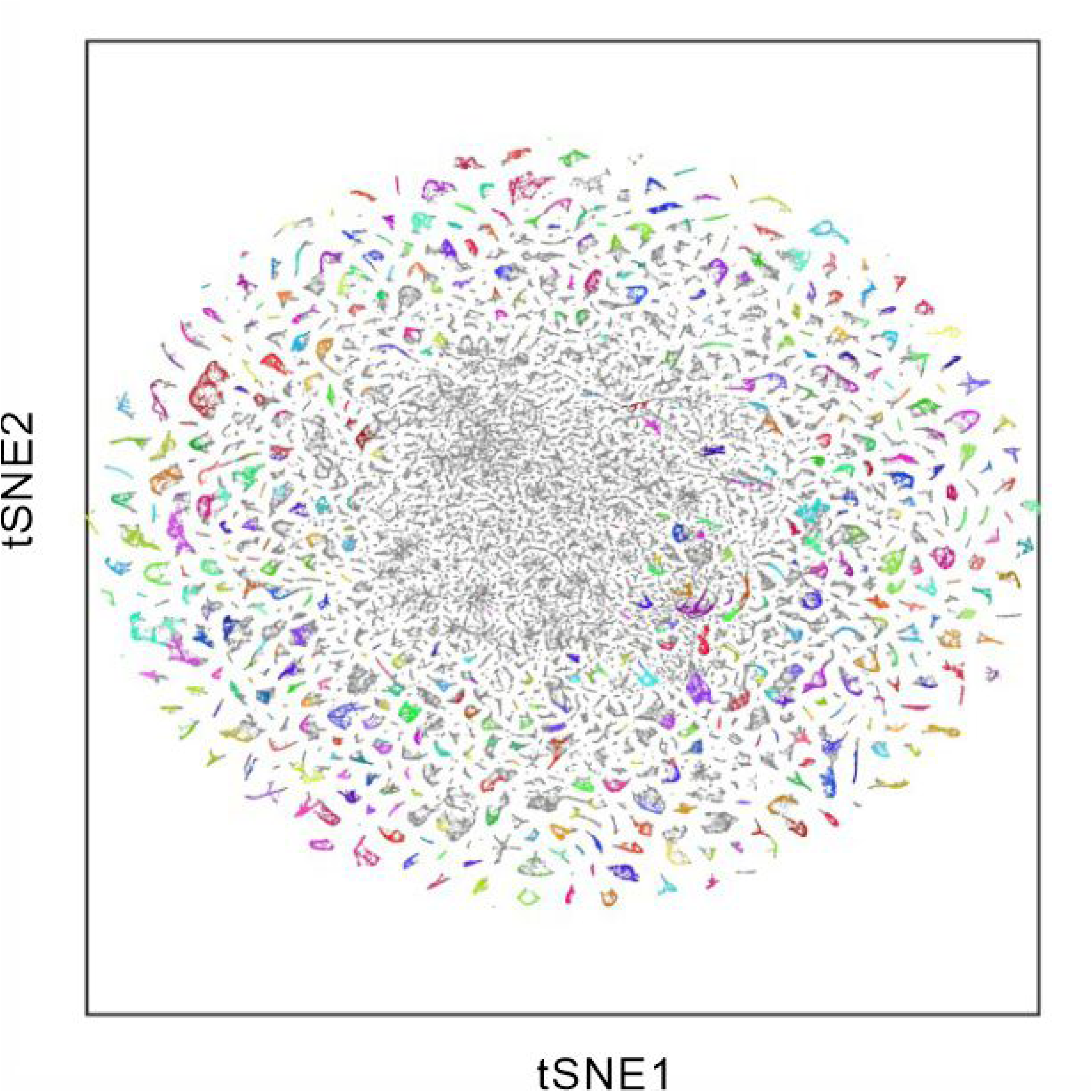
**The distribution of different neuronal components in the t-SNE map**.

**Figure S3.**
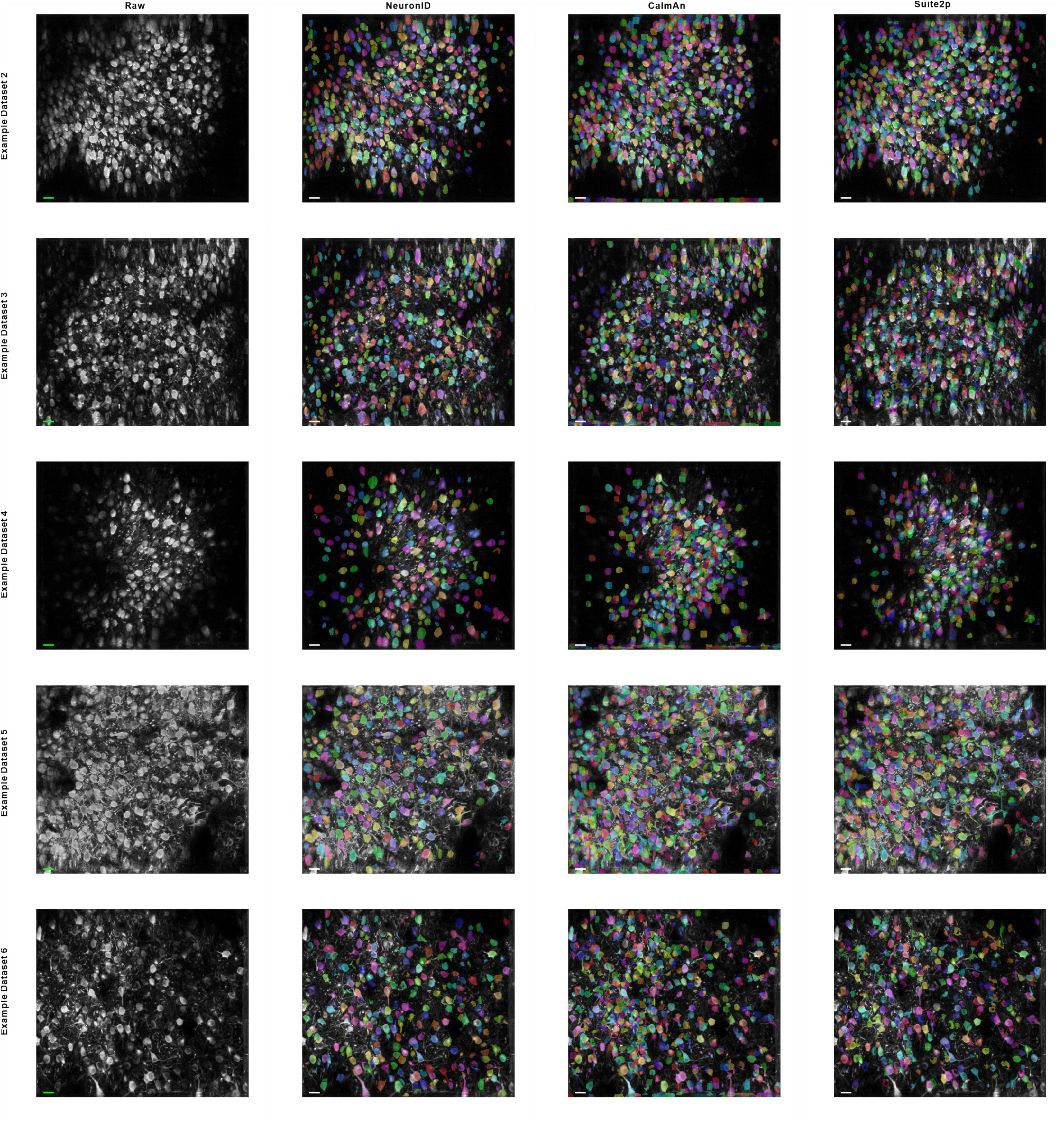
Comparative analysis among three toolkits. The max-mean projection images (Raw) and soma masks generated by NeuronID, CaImAn, and Suite2p across five example datasets. Scale bar: 10 pixels.

**Figure S4.**
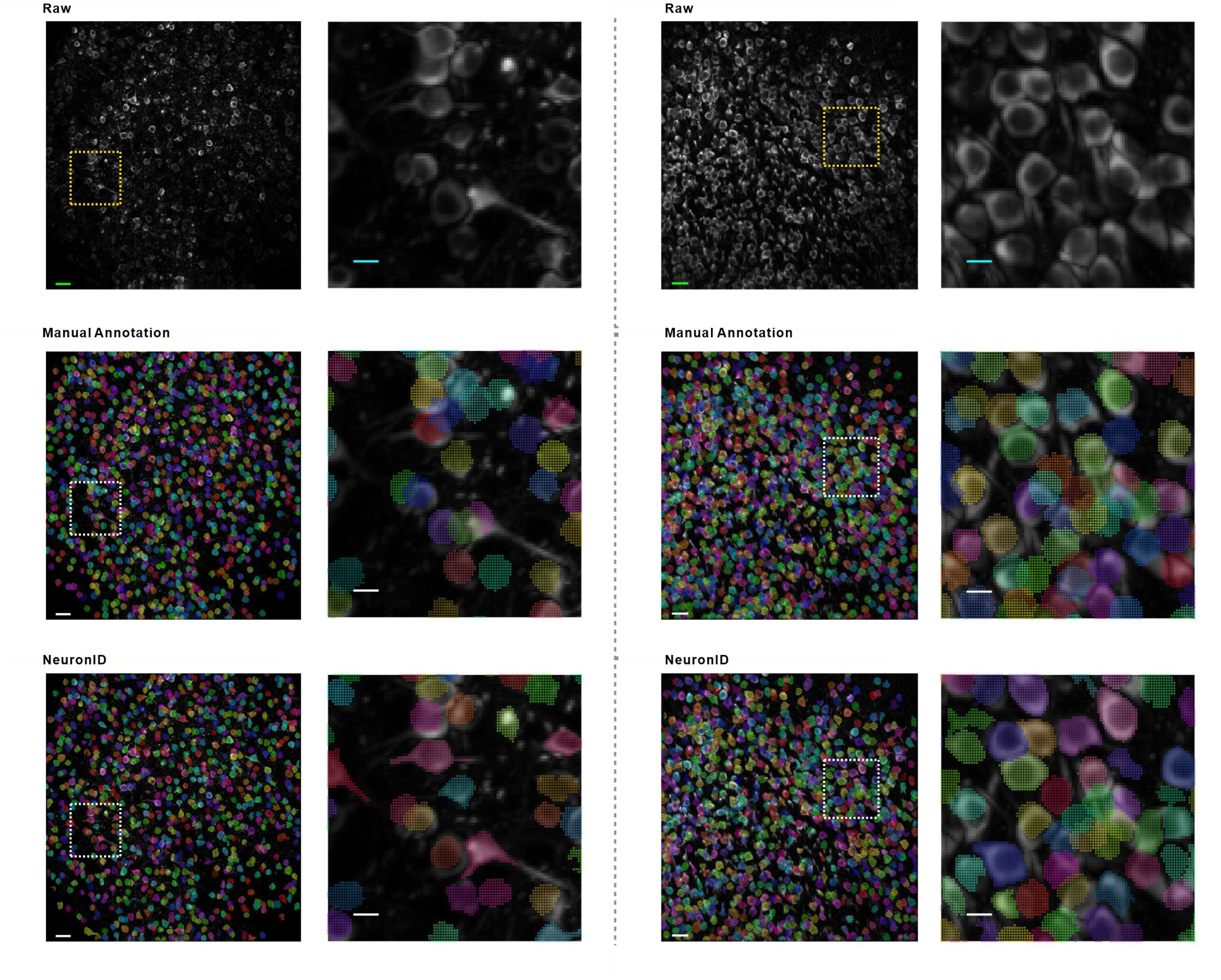
Comparative analysis of the NeuronID toolkit versus manual annotation. Max-mean projection images (Raw) and soma masks generated by the human annotation or the NeuronID toolkit. Scale bar: 10 pixels (left) or 30 pixels (right).

**Table S1.** The number of neuronal components segmented by different toolkits.

| **Dataset ID** | **NeuronID** | **CalmAn** | **Suite2p** |
| --- | --- | --- | --- |
| **Example Dataset 1** | 448 | 822 | 632 |
| **Example Dataset 2** | 375 | 689 | 440 |
| **Example Dataset 3** | 442 | 775 | 609 |
| **Example Dataset 4** | 274 | 727 | 414 |
| **Example Dataset 5** | 275 | 687 | 302 |
| **Example Dataset 6** | 241 | 632 | 258 |

## References

1. Denk, W., Strickler, J.H., and Webb, W.W. (1990). Two-Photon Laser Scanning Fluorescence Microscopy. Science 248, 73–76. 10.1126/science.2321027.

2. Grienberger, C., Giovannucci, A., Zeiger, W., and Portera-Cailliau, C. (2022). Two-photon calcium imaging of neuronal activity. Nature Reviews Methods Primers 2, 67. 10.1038/s43586-022-00147-1.

3. Chen, T.-W., Wardill, T.J., Sun, Y., Pulver, S.R., Renninger, S.L., Baohan, A., Schreiter, E.R., Kerr, R.A., Orger, M.B., Jayaraman, V., et al. (2013). Ultrasensitive fluorescent proteins for imaging neuronal activity. Nature 499, 295–300. 10.1038/nature12354.

4. Tian, L., Hires, S.A., Mao, T., Huber, D., Chiappe, M.E., Chalasani, S.H., Petreanu, L., Akerboom, J., McKinney, S.A., Schreiter, E.R., et al. (2009). Imaging neural activity in worms, flies and mice with improved GCaMP calcium indicators. Nature Methods 6, 875–881. 10.1038/nmeth.1398.

5. Inoue, M. (2021). Genetically encoded calcium indicators to probe complex brain circuit dynamics in vivo. Neuroscience Research 169, 2–8. 10.1016/j.neures.2020.05.013.

6. Miyawaki, A., Llopis, J., Heim, R., McCaffery, J.M., Adams, J.A., Ikura, M., and Tsien, R.Y. (1997). Fluorescent indicators for Ca2+based on green fluorescent proteins and calmodulin. Nature 388, 882–887. 10.1038/42264.

7. Nakai, J., Ohkura, M., and Imoto, K. (2001). A high signal-to-noise Ca2+ probe composed of a single green fluorescent protein. Nature Biotechnology 19, 137–141. 10.1038/84397.

8. Bai, L., Cong, L., Shi, Z., Zhao, Y., Zhang, Y., Lu, B., Zhang, J., Xiong, Z.-Q., Xu, N., Mu, Y., and Wang, K. (2024). Volumetric voltage imaging of neuronal populations in the mouse brain by confocal light-field microscopy. Nature Methods 21, 2160–2170. 10.1038/s41592-024-02458-5.

9. Zong, W., Wu, R., Li, M., Hu, Y., Li, Y., Li, J., Rong, H., Wu, H., Xu, Y., Lu, Y., et al. (2017). Fast high-resolution miniature two-photon microscopy for brain imaging in freely behaving mice. Nature Methods 14, 713–719. 10.1038/nmeth.4305.

10. Sofroniew, N.J., Flickinger, D., King, J., and Svoboda, K. (2016). A large field of view two-photon mesoscope with subcellular resolution for in vivo imaging. Elife 5. 10.7554/eLife.14472.

11. Helmchen, F., and Denk, W. (2005). Deep tissue two-photon microscopy. Nature Methods 2, 932–940. 10.1038/nmeth818.

12. de Vries, S.E.J., Lecoq, J.A., Buice, M.A., Groblewski, P.A., Ocker, G.K., Oliver, M., Feng, D., Cain, N., Ledochowitsch, P., Millman, D., et al. (2020). A large-scale standardized physiological survey reveals functional organization of the mouse visual cortex. Nature Neuroscience 23, 138–151. 10.1038/s41593-019-0550-9.

13. Pachitariu, M., Stringer, C., Dipoppa, M., Schröder, S., Rossi, F., Dalgleish, H., Carandini, M., and Harris, K. (2016). Suite2p: beyond 10,000 neurons with standard two-photon microscopy. bioRxiv.

14. Giovannucci, A., Friedrich, J., Gunn, P., Kalfon, J., Brown, B.L., Koay, S.A., Taxidis, J., Najafi, F., Gauthier, J.L., Zhou, P., et al. (2019). CaImAn an open source tool for scalable calcium imaging data analysis. eLife 8, e38173. 10.7554/eLife.38173.

15. Shipley, F.B., Dani, N., Xu, H., Deister, C., Cui, J., Head, J.P., Sadegh, C., Fame, R.M., Shannon, M.L., Flores, V.I., et al. (2020). Tracking Calcium Dynamics and Immune Surveillance at the Choroid Plexus Blood-Cerebrospinal Fluid Interface. Neuron 108, 623–639.e610. 10.1016/j.neuron.2020.08.024.

16. Dombeck, D.A., Khabbaz, A.N., Collman, F., Adelman, T.L., and Tank, D.W. (2007). Imaging large-scale neural activity with cellular resolution in awake, mobile mice. Neuron 56, 43–57. 10.1016/j.neuron.2007.08.003.

17. Pnevmatikakis, E.A., and Giovannucci, A. (2017). NoRMCorre: An online algorithm for piecewise rigid motion correction of calcium imaging data. Journal of Neuroscience Methods 291, 83–94. 10.1016/j.jneumeth.2017.07.031.

18. Malik, W.Q., Schummers, J., Sur, M., and Brown, E.N. (2011). Denoising two-photon calcium imaging data. PLoS One 6, e20490. 10.1371/journal.pone.0020490.

19. Lecoq, J., Oliver, M., Siegle, J.H., Orlova, N., Ledochowitsch, P., and Koch, C. (2021). Removing independent noise in systems neuroscience data using DeepInterpolation. Nature Methods 18, 1401–1408. 10.1038/s41592-021-01285-2.

20. Kerr, J.N.D., Greenberg, D., and Helmchen, F. (2005). Imaging input and output of neocortical networks in vivo. Proceedings of the National Academy of Sciences 102, 14063–14068. 10.1073/pnas.0506029102.

21. Demas, J., Manley, J., Tejera, F., Barber, K., Kim, H., Traub, F.M., Chen, B., and Vaziri, A. (2021). High-speed, cortex-wide volumetric recording of neuroactivity at cellular resolution using light beads microscopy. Nature Methods 18, 1103–1111. 10.1038/s41592-021-01239-8.

22. Park, E.-Y., Cai, X., Foiret, J., Bendjador, H., Hyun, D., Fite, B.Z., Wodnicki, R., Dahl, J.J., Boutin, R.D., and Ferrara, K.W. Fast volumetric ultrasound facilitates high-resolution 3D mapping of tissue compartments. Science Advances 9, eadg8176. 10.1126/sciadv.adg8176.

23. Sadakane, O., Masamizu, Y., Watakabe, A., Terada, S.-I., Ohtsuka, M., Takaji, M., Mizukami, H., Ozawa, K., Kawasaki, H., Matsuzaki, M., and Yamamori, T. (2015). Long-Term Two-Photon Calcium Imaging of Neuronal Populations with Subcellular Resolution in Adult Non-human Primates. Cell Reports 13, 1989–1999. 10.1016/j.celrep.2015.10.050.

24. Broussard, G.J., Liang, Y., Fridman, M., Unger, E.K., Meng, G., Xiao, X., Ji, N., Petreanu, L., and Tian, L. (2018). In vivo measurement of afferent activity with axon-specific calcium imaging. Nature Neuroscience 21, 1272–1280. 10.1038/s41593-018-0211-4.

25. Peng, J., Yang, D., Xu, C., and Xu, T. (2025). The Neural Basis of Quantity Discrimination in Mice. bioRxiv, 2025.2007.2026.666982. 10.1101/2025.07.26.666982.

26. F.R.S., K.P. (1901). LIII. On lines and planes of closest fit to systems of points in space. Philosophical Magazine Series 1 *2*, 559–572.

27. van der Maaten, L., and Hinton, G. (2008). Viualizing data using t-SNE. Journal of Machine Learning Research 9, 2579–2605.

28. Macqueen, J.J.P.S.M.S., and Probability, t. (1967). Some methods for classification and analysis of multivariate observations. Proceedings of the 5th Berkeley Symposium on Mathematical Statistics and Probability 1.

29. Pascanu, R., Mikolov, T., and Bengio, Y. (2013). On the difficulty of training recurrent neural networks.

30. Scheenen, W.J.M., Makings, L.R., Gross, L.R., Pozzan, T., and Tsien, R.Y. (1996). Photodegradation of indo-1 and its effect on apparent Ca2+ concentrations. Chemistry & Biology 3, 765–774. 10.1016/S1074-5521(96)90253-7.

31. Komiyama, T., Sato, T.R., O’Connor, D.H., Zhang, Y.-X., Huber, D., Hooks, B.M., Gabitto, M., and Svoboda, K. (2010). Learning-related fine-scale specificity imaged in motor cortex circuits of behaving mice. Nature 464, 1182–1186. 10.1038/nature08897.

32. Christen, P., Hand, D.J., and Kirielle, N. (2023). A Review of the F-Measure: Its History, Properties, Criticism, and Alternatives. ACM Computing Surveys 56, Article 73. 10.1145/3606367.

